# EPSTEIN-BARR VIRUS LMP1 ENHANCES LEVELS OF MICROVESICLE-ASSOCIATED PD-L1

**DOI:** 10.1101/2022.01.14.476429

**Authors:** Monica Abou Harb, Li Sun, David G. Meckes

## Abstract

Extracellular vesicles (EVs) circulate throughout the body and carry cargo that can be conferred to proximal or distant cells, making them major delivery vehicles for cellular communication. Epstein-Barr virus (EBV) infected cells release EVs that contain viral proteins such as the major viral oncogene, latent membrane protein 1 (LMP1). LMP1 has been shown to regulate the cellular gene expression of programmed cell death protein 1 ligand (PD-L1). PD-L1, a protein that suppresses the immune system by binding to PD-1, (a receptor found on cytotoxic T cells). PD-L1 has been recently found to be packaged into small EVs contributing to immune evasion of lung cancer cells. Recent studies establish that MVs are shed in very large amounts by tumor cells, and that elevated levels of MVs correlate to disease metastasis and cancers being more aggressive. Here, we demonstrate PD-L1 enrichment in MVs released from nasopharyngeal carcinoma cells and an important function of EBV LMP1 in regulating PD-L1 levels in MVs. These PD-L1+ MVs containing LMP1 likely contribute to the immunosuppressive microenvironment found in EBV-associated cancers.

**Importance:** Accumulating evidence over the past decade supports that viruses utilize EVs and associated pathways to incorporate viral products to evade eliciting an immune response, while concurrently enabling viral spread or persistence within the host. Considering that viral proteins confer very strong antigenic peptides that will be recognized by T cells, the regulation of the PD-1 pathway by the overexpression of MV-associated PD-L1 may be a strong immune evasion tactic utilized by viruses. The discovery that EBV LMP1 increases PD-L1 microvesicle secretion, identifies a new therapeutic target in immune blockade therapy. We expect that our findings will begin to clarify the mechanism of LMP1-mediated enhanced packaging of PD-L1 into MVs and may produce more specific targets to treat EBV-associated cancers. Consequently, identifying whether a disease is of viral origin through predictive MV biomarkers could further allow for more targeted therapies.

## Introduction

Epstein-Barr virus (EBV) is a human herpesvirus that persistently infects over 90% of the world’s population (1). Most individuals are asymptomatic; however, in immunocompromised or genetically susceptible individuals, EBV infection can cause various lymphomas and carcinomas (1). PD-L1 normally functions by suppressing an individual’s immune system from attacking their own cells within the body. Any pathogenic, viral, or foreign substance trying to invade the body is combated by the immune system; however, certain cancer cells take advantage of the functions of PD-L1 and overexpress this gene in order to escape immune detection and maintain proliferation and survival. PD-L1 exists in the blood in two different forms: extracellular vesicle-associated or a soluble protein. One of the first observations of PD-L1 being expressed on small extracellular vesicles (EVs) was in 2011 where investigators detected exosomal PD-L1 extracted from human samples of plasma and urine (2). Additionally, PD-L1 has been found on the surface of antigen presenting cells such as B cells, mast cells, and certain epithelial and endothelial cells (3, 4). It is highly expressed on the surface of tumor cells, T lymphocytes and tumor associated macrophages (5). Additionally, it can be found in many tissues such as muscles and nerves (6). PD-L1 suppresses the immune system by binding to PD-1, a receptor found on cytotoxic T cells. The PD-1 pathway involves several interactions leading to the inhibition of T-regulatory functions and the production of interleukins which results in a reduced immune response, T cell apoptosis and exhaustion, and the suppression of dendritic cells (7). These cumulative immunosuppressive effects create a very strong protective barrier for cancerous cells to grow uncontrollably and eventually metastasize throughout the body. PD-L1 may also play an important role in immune evasion strategies adopted by viruses and other human pathogens.

Some of the more recent data has suggested an important role for the PD-L1 ligand as a mechanism of achieving an immune resistant tumor microenvironment and establishing latency within the host in EBV infection. Expression of PD-L1 is significantly higher in EBV positive cells compared to EBV negative cells (8). It has been shown that high EBV copy number per infected cell is correlated with PD-L1 expression such as in EBV associated gastric carcinoma (9). The presence of high levels of PD-1 and PD-L1 in EBV+ patients is directly linked to the development of high-risk Hodgkin lymphoma (10). Considering that LMP1, (major viral oncogene expressed in EBV associated cancers), has been established in regulating PD-L1 levels within the cell and manipulates EV cargo and functions, we hypothesized that LMP1 may influence PD-L1 EV levels as an immune evasion mechanism. Ultimately, this would result in an immunosuppressive tumor microenvironment favoring viral sustainability and propagating infection, not only to neighboring cells, but cells throughout the body as well.

Viruses share many similarities with EVs, such as the transfer of functional cellular proteins, RNAs, and miRNAs into neighboring cells (11). The formation of large EVs or microvesicles (MVs) involves transport of cargo to the plasma membrane, lipid membrane redistribution, budding, and finally a form of vesicle pinching upon reaching the membrane which leads to the release of the vesicle (12). Shed MVs are distinct in size, cargo, and mechanism of formation from the other subpopulation of EVs known as small EVs. Both small EVs and MVs encapsulate and transfer cargo including proteins, miRNAs, and RNA transcripts to other cells. Caveolin-1 is essential for sorting of selected miRNAs into MVs (13). Both viruses and MVs emerge from the plasma membrane at lipid raft organizing centers indicating a similar form of biogenesis (14). Elevated levels of EVs in tumor microenvironments and within the circulation correlate to a more advanced disease stage in cancers. MVs play a role in acting as a bridge of cellular communication between cancer and the surrounding stroma to alter the surrounding tumor environment in a way that enhances their growth and survival. MVs shed from a highly metastatic melanoma cell line, were able to confer metastatic characteristics to poorly metastatic cells (15). Recent data has indicated a role for PD-L1 expression in small EVs; however, little is known about PD-L1 expression in MVs, the mechanism that leads to its upregulation, and the localization of PD-L1 in these different EV subpopulations.

Previous studies have already implicated a role for MVs in immune suppression. For example, MVs derived from colorectal carcinoma or human melanoma, suppressed monocyte differentiation into antigen presenting cells and suppressed T lymphocytes through TGF-β, both *in vivo* and *in vitro* (16). LMP1, (a major viral oncoprotein present in EVs), has been shown to be involved in indirectly inhibiting humoral immune response(17, 18). LMP1 has a major role in the lifecycle of EBV, is an important contributor to many EBV-associated cancers, and is packaged and released from infected cells into small EVs (19). LMP1 activates several signaling pathways such as the JAK/STAT, MAPK, and NF-KB pathways which have been shown to upregulate PD-L1 expression as well and are known to be commonly constitutively activated in a multitude of cancers (19). Here we demonstrate the enrichment of PD-L1 in MVs in response to levels of LMP1 and clarify localization, co-immunoprecipitation, and the regulation of these MVs in response to LMP1 levels.

## Results

### PD-L1 levels are elevated in MVs from NPC cells with LMP1 expression

EBV infected cells have different expression of viral transcriptional programs referred to as latency I, II, and III. EBV Latency II, was originally identified in biopsies of NPC, which was characterized by the co-expression of LMP1, LMP2, and EBNA1 transcripts (20). NPC tissues expressing LMP1 are significantly associated with poor overall survival in NPC patients and lymph node metastasis (21, 22). We decided to investigate two different NPC cell lines with varying levels of LMP1. In the first model, an EBV-negative nasopharyngeal carcinoma cell line (HK1) was compared to HK1 containing a GFP-tagged LMP1 tetracycline-inducible system (HK1 LMP1). The HK1 LMP1 cells were treated with doxycycline to activate LMP1 expression (Fig. 1 A). Additionally, to investigate PD-L1 expression in the context of lower stable expression of LMP1, EBV-negative nasopharyngeal carcinoma (HNE-1) cells were used. HNE-1 cells were transduced with a vector control (pBabe) or hemagglutinin (HA)-tagged LMP1 retroviral vector (HNE-1 pBabe LMP1) that constitutively expresses low levels of LMP1 (Fig. 1 B). LMP1 expression was detectable in the HNE-1 pBabe LMP1 but notably lower than the HK1 LMP1 inducible model. The lower levels of LMP1 have been previously reported to closely exhibit expression patterns found in EBV-infected cell lines (23). We harvested the cell lysates, apoptotic bodies (2K), microvesicles (10K), and small EVs (100K), and analyzed them by immunoblot (24). The results showed an increase in PD-L1 levels with elevated levels of LMP1, with PD-L1 noticeably enriched within the MV subpopulation of EVs compared to small EVs and apoptotic bodies (Fig 1.A and B). Quantification of the levels of LMP1 and PD-L1 specifically in the MV subpopulation normalized to levels of (HK1 and HNE-1) showed a significant increase in PD-L1 and LMP1 in the MVs of the NPC cell lines expressing LMP1 compared to the wild type cell lines with no LMP1 (Fig. 1C and D). Quantification also demonstrated higher PD-L1 expression in the MVs compared to the other two subpopulations of EVs as seen in (Fig. 1 E and F). In conclusion, these results further support the correlation between LMP1 expression and increased PD-L1 being released from the cell in association with MVs.

**Fig 1.**
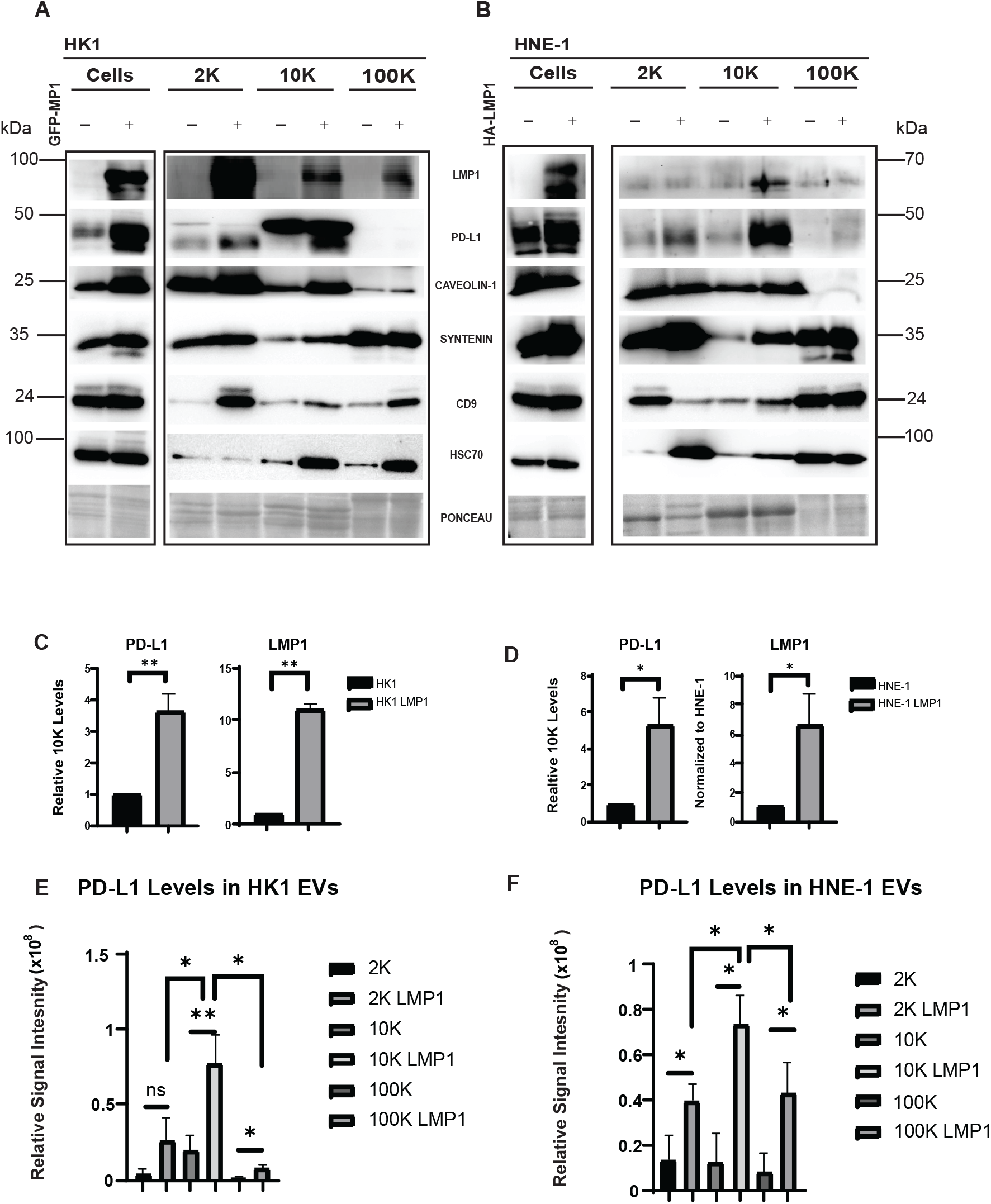
LMP1 expression in NPC cells leads to enhanced PD-L1 levels in MVs. Cell lysates, 2K, 10K, and 100K were loaded at equal protein concentration of 20μg. (A and B) Immunoblot analyses of HK1 GFP-tagged LMP1 and HNE-1 HA-tagged LMP1 demonstrate increased PD-L1 expression in the MV subpopulation compared to HK1 and HNE-1, respectively. (C and D) Quantification of 3 western blots from independent experiments demonstrate significantly higher levels of PD-L1 expression in MVs expressing LMP1 compared to control cell lines HK1 and HNE-1. (E and F) Quantification of 3 western blots in NPC cells expressing LMP1 revealed significantly higher levels of PD-L1 in the MVs compared to the other two EV subpopulations. **, P<0.01; *, P<0.05.

### PD-L1 is upregulated at the transcriptional level in LMP1 expressing cells and detected in MVs

To further understand whether PD-L1 is also being impacted at an mRNA level due to LMP1 expression, RT-qPCR was performed on cells and MVs from both HK1 wild type cells and HK1 LMP1 induced cells. HK1 LMP1 containing cells significantly increased the expression of PD-L1 and EV biogenesis related genes (CD9, CD63, CD81, Sytenein-1, and HRS) compared to HK1 cells (Fig. 2A). Interestingly, LMP1 expression did not influence the mRNA levels of PD-L1 in MVs but did have significantly higher levels of CD9, CD63, CD81, Syntenin-1, and HRS mRNA when compared to HK1 (Fig. 2B). CD9, CD63 and CD81 are enodosome-specific tetraspanins that are enriched in exosome (samall EV) membranes. CD63 has been found to regulate EBV LMP1 exosomal packaging (18) and HRS and Syntenin-1 have previously been shown to induce EV formation through EBV LMP1 (25). Altogether, this data suggests that PD-L1 is upregulated at the transcriptional level in LMP1 expressing cells but the level of mRNA transcripts in the MVs remain unchanged.

**Fig 2.**
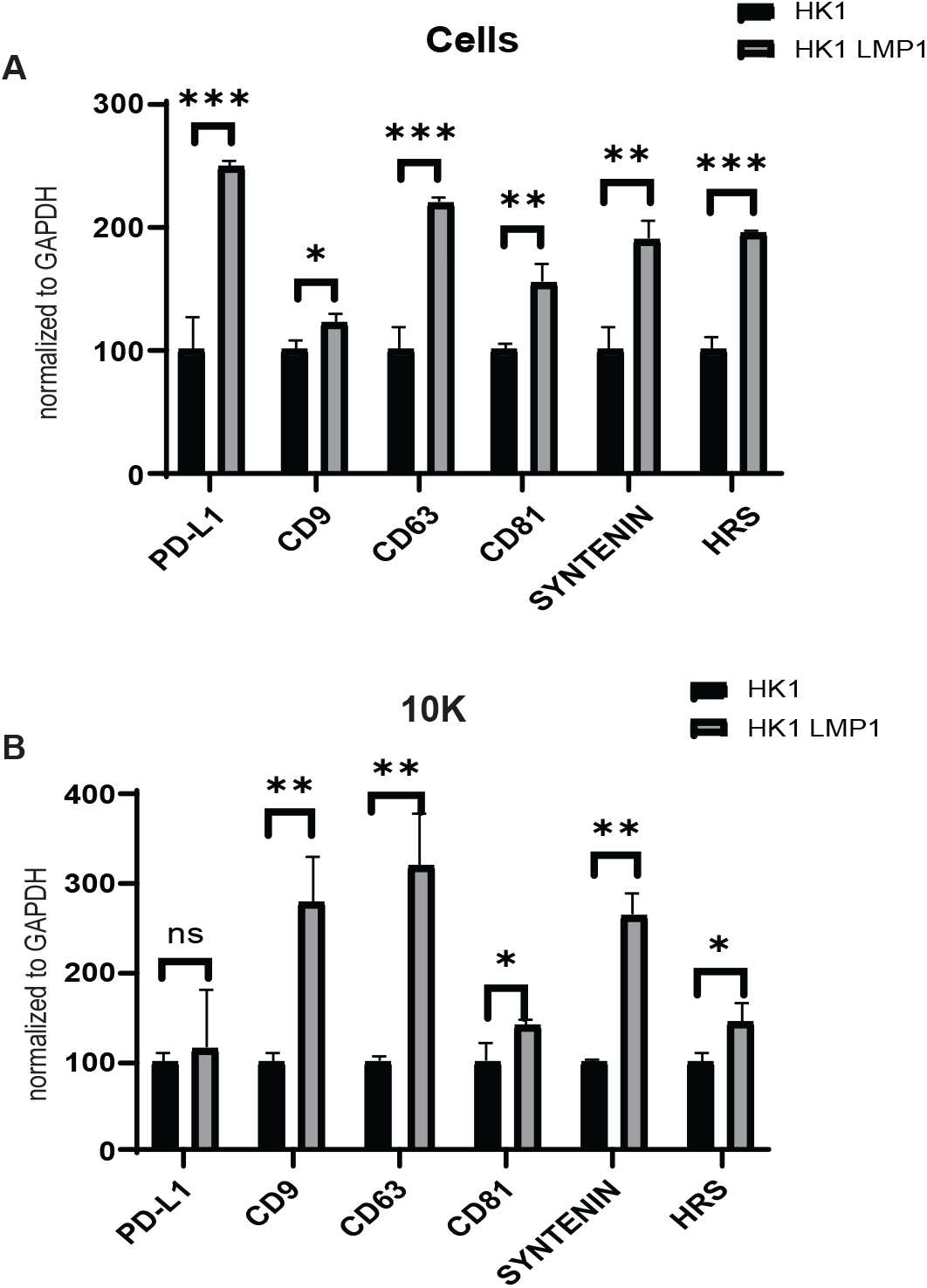
PD-L1 is upregulated at the transcriptional level in LMP1 expressing cells and is detected in MVs. RT-qPCR result of mRNA from normal HK1 cells or HK1 LMP1 induced cells (A) and MVs (B). The results were normalized to GAPDH. ***, P<0.001 **, P<0.01; *, P<0.05.

### Characterization of MVs from NPC cells

MVs are characterized as being within the 100 to few micrometers size range (26). In order to confirm size range of these MVs harvested from NPC cells with or without LMP1, Transmission Electron microscopy (TEM) imaging was performed. Both the HK1 and HK1 GFP LMP1 inducible MV subpopulations were visualized to be around 200nm and exhibited similar phenotypes under the microscope, as seen in (Fig. 3A and B). Additionally, nanoparticle tracking was performed to further validate the EM results and to identify differences in MVs harvested from NPC cells expressing LMP1 compared to wild type NPC cells. The tracking data in (Fig. 3C) revealed that there was a significant increase in particles/ml for the MVs harvested from the high expressing LMP1 NPC cell line compared to the wild type cell line, however, there was no significant difference in particles/cell between the two cell lines (Fig. 3D). This is different than what was previously observed for enhancement of small EV production due to LMP1 expression (27). The reasoning to why increased particles per ml is observed could be due to the fact that NPC cells expressing LMP1 grow at a faster rate than the wild type cells resulting in more MVs being released per ml of media. Both the mean and mode size of the MVs secreted from the high LMP1 expressing cell line was significantly increased compared to wild type MVs (Fig. 3E and F). This led us to question whether these larger secreted MVs carry more protein cargo. The tracking and protein analyses revealed that the MVs from the high expressing LMP1 cell line had a significant increase in total protein content compared to the wild type MVs (Fig. 3F). The nanoparticle tracking data suggests that there are not more MVs secreted per cell in the high expressing LMP1 cells but there is more protein per MV. Therefore, cells expressing LMP1 secrete larger MVs with increased protein content that may alter the immune response within the infected microenvironment.

**Fig 3.**
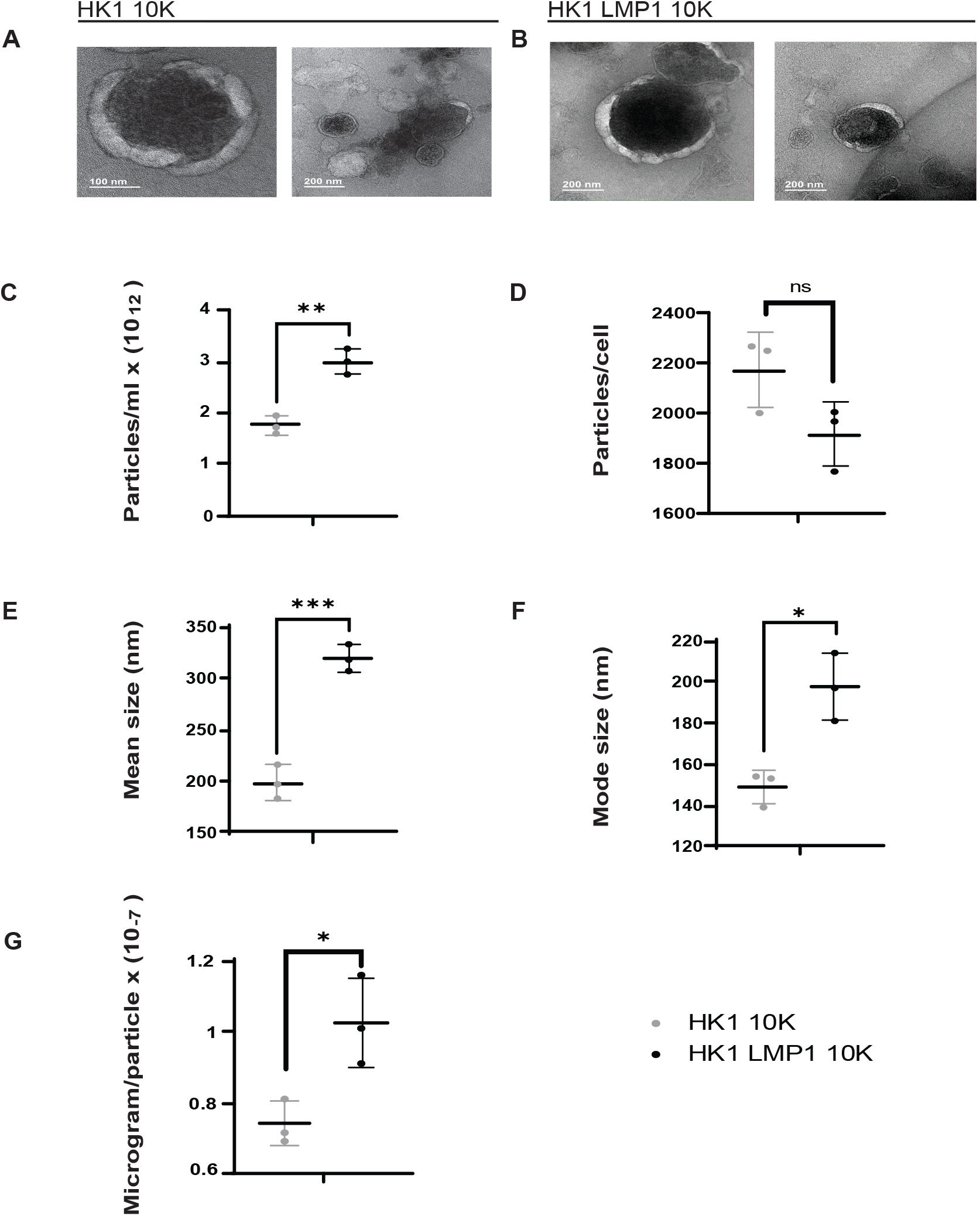
Characterization of MVs from NPC cells. (A and B) MVs harvested by 10,000 RPM spin from HK1 and HK1 LMP1 induced cells were examined by electron microscopy. (C-G) Nanoparticle tracking analysis showed significant increase in particles/ml, microgram/particle, mean, and mode size but not in particles/cell from MVs harvested from HK1 LMP1 cells compared to HK1 control cells. ***, P<0.001 **, P<0.01; *, P<0.05.

### PD-L1 is enhanced on the surface of MVs due to LMP1

PD-L1 has previously been reported to be expressed on the surface of small EVs similar to cell surface PD-L1 membrane topology, in which the extracellular domain of PD-L1 is surface exposed (28). PD-L1 extracellular exposure is required for it to exert its immunosuppressive effects, since PD-L1 needs to be exposed on the surface to bind to its receptor PD-1 which is found on cytotoxic T cells. Flow cytometry on MVs from HK1 wild type cells and HK1 LMP1 induced cells expressing LMP1 was done. The MVs from both cell lines were incubated with surface PD-L1 antibody or isotype control. Flow cytometry analyses revealed elevated surface PD-L1 expression in the MVs from HK1 GFP tagged LMP1 (green) compared to the MVs from the HK1 wild type cells (blue) (Fig. 4A). In order to further understand PD-L1 localization on these MVs, scanning transmission electron microscopy (STEM) was undertaken to observe the MVs after immunogold labelling of PD-L1 with antibodies that specifically bind to PD-L1 on the extracellular domain. Upon examination of both the prepared EM grids, gold labelling of PD-L1 was only observed on the HK1 LMP1 MVs (Fig. 4B and C). Quantification of gold particles per MV revealed a significant increase in gold particles in the HK1 LMP1 MVs compared to the wild type MVs (Fig. 4D). Research has shown PD-L1 localization to be on the surface of small EVs; however, more current studies have found PD-L1 to be present on both the surface and within small EVs (29). To further understand PD-L1 localization within the MV, a protease protection experiment was done using two different PD-L1 antibodies. For this experiment, Syntenin and CD71 were used as the positive and negative controls, respectively. Syntenin is a cytosolic adaptor located within EVs that binds to the intracellular domain (ICD) of syndecans (30), and CD71 is a transferrin receptor found on the surface of EVs. In (Fig. 4E), a PD-L1 antibody that recognizes endogenous levels of PD-L1 showed a decrease in band size upon the addition of trypsin. In (Fig. 4F) a PD-L1 antibody that recognizes the extracellular domain of PD-L1 was used, and upon trypsin addition little PD-L1 is observed. This reveals that there is some PD-L1 surface exposure on these MVs secreted from high LMP1 expressing cells; however, some PD-L1 may also be present inside MVs or is resistant to Trypsin cleavage. Taken together, LMP1 seems to play a role in increasing PD-L1 exposure to the surface of MVs which may influence the immunosuppressive effects of MVs released from EBV infected cells.

**Fig 4.**
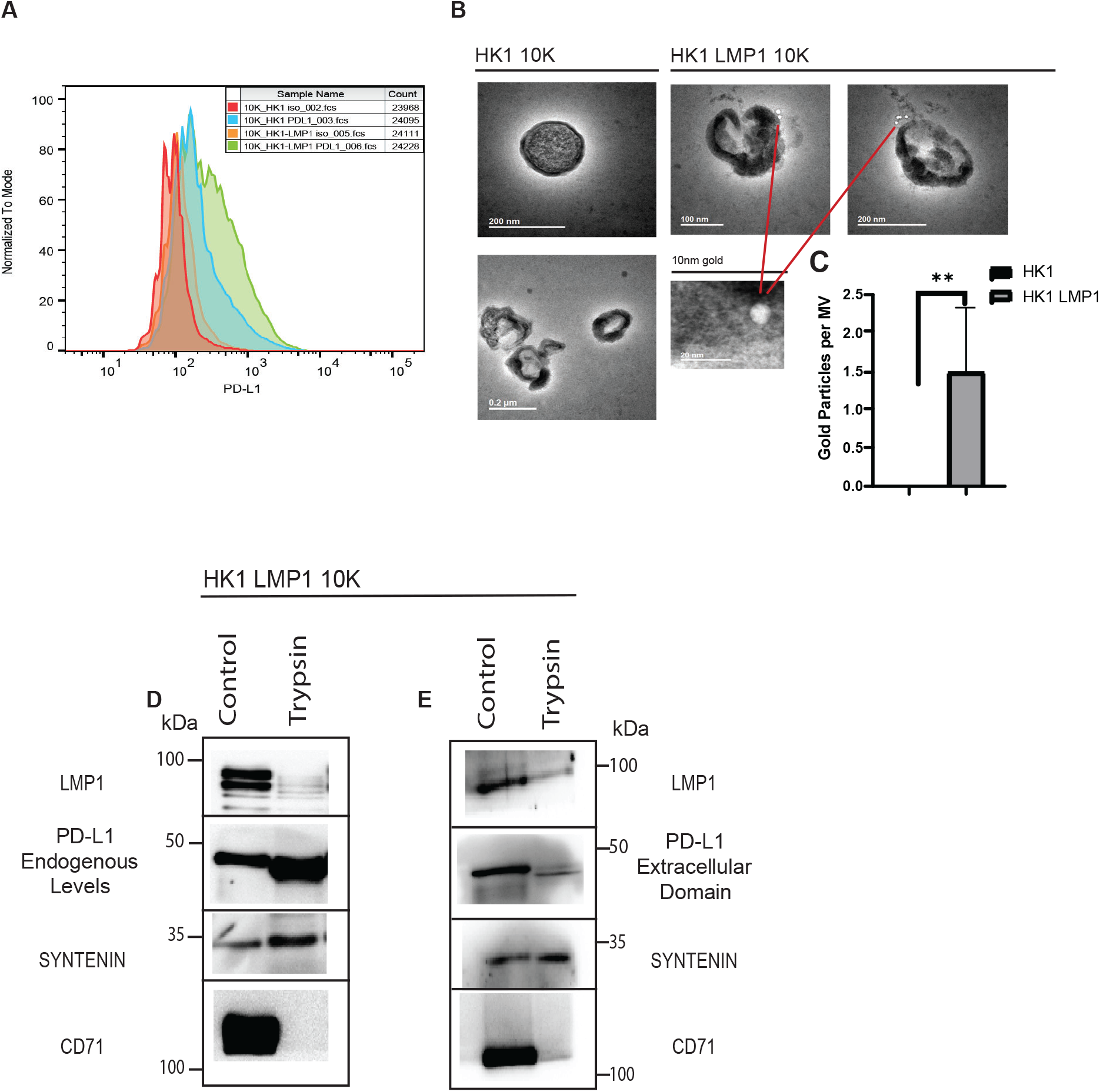
LMP1 enhances PD-L1 MV-associated surface levels. (A) Flow cytometry analysis on MVs harvested from HK1 and HK1 LMP1 induced cells stained for either isotype control or PD-L1 surface antibody. (B and C) PD-L1 immunogold labelling of HK1 and HK1 LMP1 MVs. (D) Quantification of gold particles per MV from 10 HK1 and HK1 LMP1 induced MVs. (E and F) Immunoblot analyses of HK1 LMP1 MVs with and without trypsin. Different PD-L1 antibodies were used to probe for endogenous levels of PD-L1 (E) and extracellular domain specific PD-L1 (F). CD71 and Syntenin-1 are used as negative and positive controls, respectively. **, P<0.01.

### LMP1 and PD-L1 localize within the same MV subpopulation

LMP1 and PD-L1 have previously been demonstrated to be expressed on small EVs (28, 31). Here, we aimed to investigate whether LMP1 was present in PD-L1 positive MVs following LMP1 induction of NPC cells. First, Optiprep density gradient purification of MVs was performed. This technique uses iodixanol and ultra-centrifugal force to further purify EV subpopulations into fractions based on their buoyant density (24). This can be used to determine the exact density and fractions within the MV pellet that contain PD-L1 and whether it co-migrates with LMP1 in the gradient. MVs were harvested from doxycycline-treated HK1 GFP-LMP1 tetracycline-inducible cells and separated by density. The vesicles were purified on an iodixanol density gradient to separate larger MV subpopulation according to the method of Kowal et al (24). Two major populations of vesicles were observed in fractions 3 (F3) and 5 (F5) of the gradients, corresponding to densities 1.115 g/mL (third fraction = F3) and 1.145 g/mL (fifth fraction = F5) of iodixanol. MV subpopulations, F3 and F5 have previously been determined to be enriched with specific proteins. F3 usually contains unique ribosome and proteasome proteins with F5 containing more mitochondrial and ER proteins (24). In these experiments, F3 and F5 MV subpopulation of EVs were enriched for protein markers LMP1, PD-L1, and Caveolin-1 (Fig. 5A). LMP1 is present in both small EVs and larger shed microvesicles (31, 32). Furthermore, we demonstrate the co-migration of LMP1 and PD-L1 in fractions 3, 4 and 5. Fractions 3 and 5 were enriched with LMP1, PD-L1 and Caveolin-1 more so than the other fractions indicating that both LMP1 and PD-L1 co-migrate within the same density fractions in MVs. If LMP1 is present in PD-L1 positive MVs, then MVs immuno-precipitated with PD-L1 specific antibodies should contain LMP1. To test this, MVs were harvested from cells expressing LMP1 and recovered by the addition of anti-PD-L1 antibody. Ten percent of the MV input sample was used to compare to the PD-L1 immuno-precipitated MVs. Immunoblot analysis further revealed LMP1 to be present in the PD-L1 pulldown (Fig. 5B). EGFR was also pulled down in these PD-L1 positive MVs which was interesting since it has been recently demonstrated that EGFR activity can control PD-L1 surface presentation in breast cancer cells (33). Together, this data suggests that LMP1 and PD-L1 are likely in the same MV population (mostly in fractions 3 and 5) and are localized to the same vesicle within MVs.

**Fig 5.**
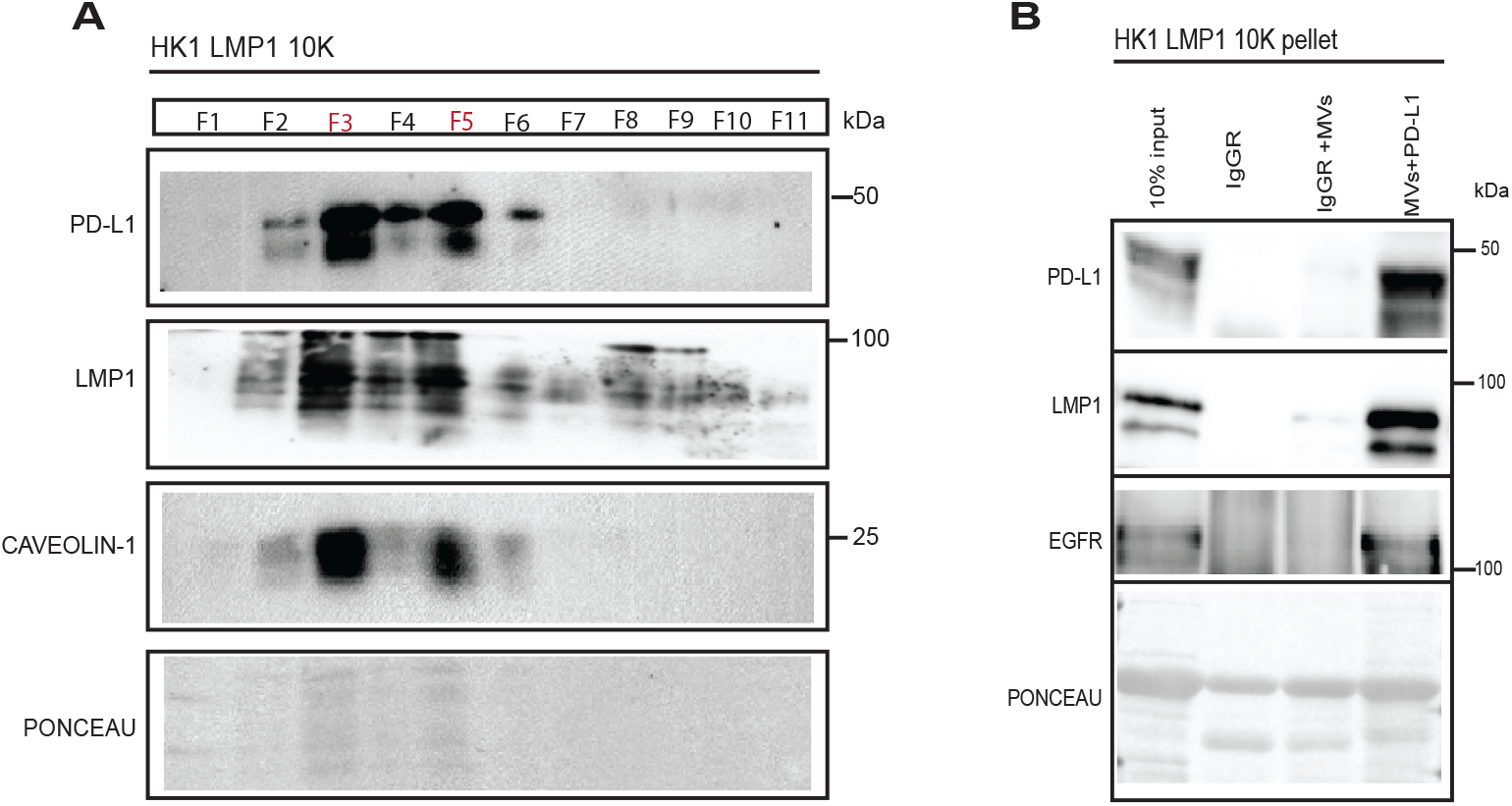
LMP1 and PD-L1 localize within the same MV subpopulation. (A) Immunoblot of iodixanol density gradient fractions of HK1 LMP1 MVs demonstrating the presence of two distinct populations enriched in LMP1, PD-L1, and Caveolin-1 corresponding to fractions (3 and 5), equal volumes loaded. (B) Immunoprecipitation of PD-L1, LMP1, and EGFR in HK1 LMP1 MVs. Equal volumes of control Rabbit IgG antibody, control rabbit IgG antibody and HK1 LMP1 MVs, and anti-PD-L1 antibody and HK1 LMP1 MVs were loaded. The first lane is 10% of the HK1 LMP1 MV sample loaded for comparison.

### LMP1 and PD-L1 colocalize in NPC cells and are enriched in lipid rafts

NPC cells have enhanced levels of PD-L1 fluorescence expression in cells that are EBV+ compared to EBV- (34). Based on the above, it was hypothesized that LMP1 might be responsible for this observation possibly even altering PD-L1 subcellular localization. Similarly, the results from (Fig. 6A-E) demonstrate enhanced PD-L1 expression and perinuclear colocalization with LMP1 in two different NPC cell lines (HK1 and HNE-1). Maximum intensity orthogonal projections of Z-stack colocalization images demonstrate PD-L1 and LMP1 perinuclear localization in live and fixed cell images (Fig. 6A and C). HNE-1 cells were transfected with RFP-tagged LMP1 and GFP-tagged PD-L1 and were examined by live cell confocal microscopy. The cells exhibited a distinct punctate perinuclear signal for LMP1 and PD-L1 (Fig. 6A). To confirm these results with endogenous levels of PD-L1 expression, and to understand what LMP1 does to PD-L1 localization within the cell, fixed cell imaging was done for HK1 LMP1 cells that were induced and un-induced with doxycycline. As seen in (Fig. 6B), PD-L1 is mostly localized perinuclearly when LMP1 is present. It seems to aggregate to one side of the cell in accordance to where LMP1 is also being accumulated. However, in the un-induced images lacking LMP1, PD-L1 was found to be mostly dispersed throughout the cell rather than localized within a sub-region like when LMP1 is present (Fig. 6C). Pearson’s correlation coefficient between LMP1 and PD-L1 in the HNE-1 transfected cells and the induced HK1 LMP1 cells, was around 0.7 and 0.8 respectively, indicating a strong correlation between these two variables (Fig. 6B and D). Additionally, CTCF values were calculated for HK1 LMP1 cells with and without doxycycline and showed very high significance in levels of PD-L1 fluorescence in the doxycycline induced HK1 LMP1 cells compared to the un-induced (Fig. 6E).

**Fig 6.**
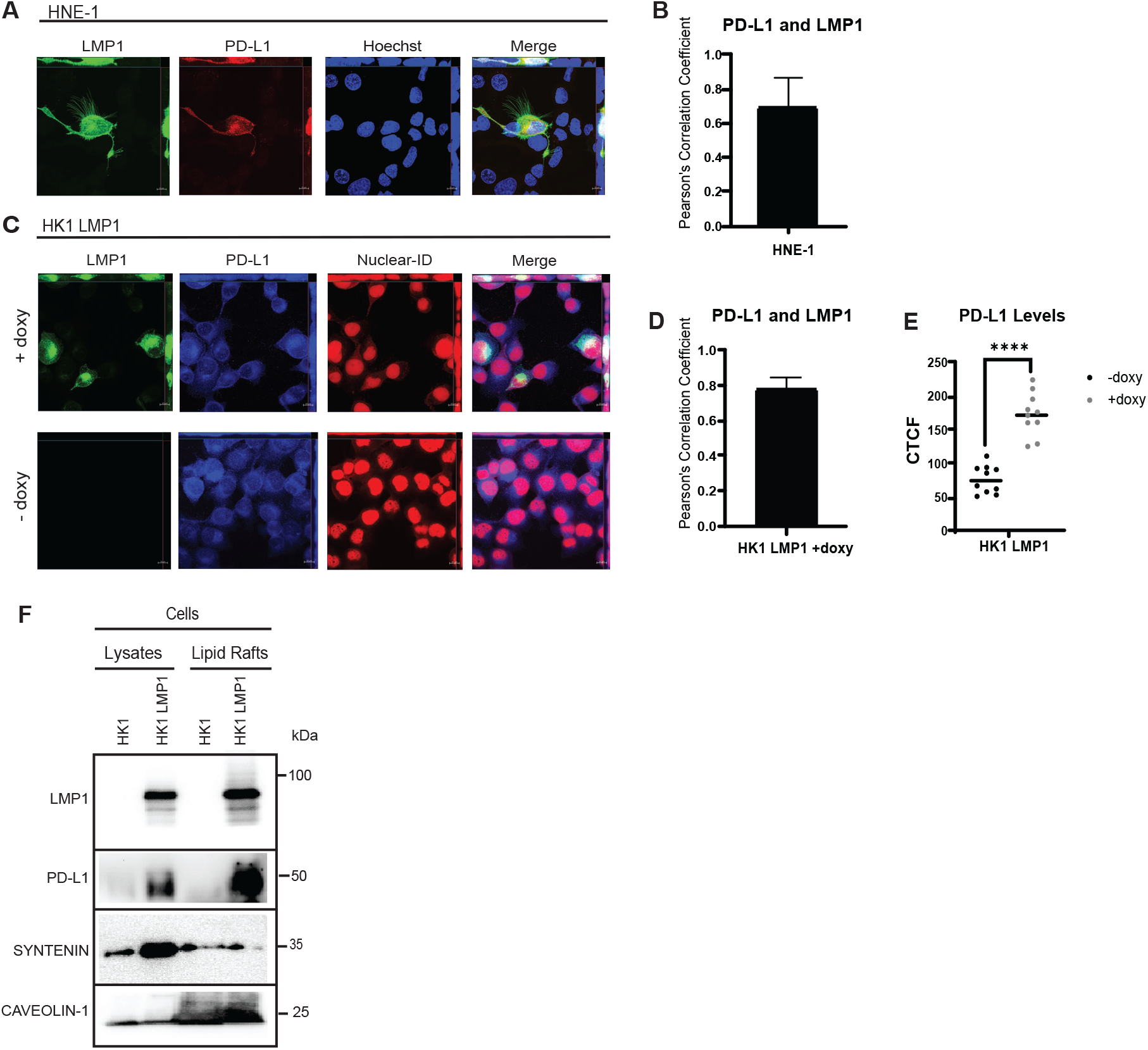
LMP1 and PD-L1 colocalize in NPC cells and are enriched in lipid rafts. All confocal images are Maximum Intensity Projections of Z-stacks, Scale bar: 10μm. (A) HNE-1 cells were transfected with GFP-tagged PD-L1, RFP-tagged LMP1, and Hoechst nuclear stain, for live-cell confocal microscopy. (C) HK1 GFP-tagged LMP1 cells induced or not induced with doxycycline were fixed, stained for PD-L1 (Alexa Fluor plus 405) and Nuclear-ID Red DNA stain and imaged by confocal microcopy. (B and D) Pearson’s Correlation Coefficient for PD-L1 and LMP1 in the HNE-1 transfected cells and induced HK1 LMP1 cells was measured in 10 cells with a value of around 0.7 and 0.8, respectively. (E) Induced HK1 LMP1 cells demonstrate significantly higher PD-L1 fluorescence compared to un-induced cells. Immunoblot analyses of whole cell lysates or lipid raft fraction of induced HK1 LMP1 cells or HK1 control cells demonstrate lipid raft enrichment of LMP1, PD-L1, and Caveolin-1. ****, P<0.0001. CTCF=corrected total cell fluorescence.

LMP1 accumulates in lipid raft microdomains within the cell to engage signaling pathways (35). Lipid rafts are distinct lipid domains enriched in cholesterol and glycosphingolipids that are involved in many roles which include but are not limited to signal transduction, membrane trafficking and the lateral compartmentalization of molecules at the cell surface (35, 36). Both viruses and MVs emerge from the plasma membrane at lipid raft organizing centers indicating a similar mechanism of biogenesis (14). The formation of MVs involves the transport of specific cargo to the plasma membrane, lipid membrane redistribution and finally a form of vesicle pinching off upon reaching the membrane which leads to the release of the vesicle (12). To test whether PD-L1 also accumulates in membrane microdomains, lipid rafts were isolated from wild type HK1 or HK1 LMP1 cells following induction of LMP1 based on their detergent-resistant biochemical properties and analyzed by immunoblot. In both cell lines raft isolates were enriched in lipid raft scaffolding protein Caveolin-1 (37). The adaptor protein Syntenin-1 was probed for since it is required for efficient LMP1 EV incorporation (25). As seen in (Fig. 6F), LMP1 and PD-L1 are enriched in lipid rafts; however, very little PD-L1 is present in wild type HK1 cells or rafts lacking LMP1. Based on this data, we conclude that PD-L1 is not only enhanced by LMP1 at the cellular level but also shifts PD-L1 localization from dispersed throughout the cell to colocalizing with LMP1 at perinuclear regions within the cell; additionally, PD-L1 and LMP1 may share a similar lipid raft associated mechanism of MV incorporation.

## Discussion

In this study, we examined the impact of EBV oncoprotein LMP1 on MV associated PD-L1 levels using a number of techniques. LMP1 has previously been found to contribute to immune evasion strategies and protect virus infected cells through the activation and upregulation of multiple immune-associated proteins (38). Moreover, LMP1 modified EVs are found to contribute to tumor progression through shaping the tumor microenvironment by enhancing tumor cell growth, migration, invasion, and suppressing the anti-tumor host response (39). LMP1 from infected cancer cells accumulate in small EVs (40). Numerous observations indicate that full-length LMP1 protein is secreted from the cell in larger EVs shed from the cell surface called microvesicles (MVs) (41). The release of LMP1 from the cell through EV pathways has been suggested as a method of escape from degradation and cell-to-cell transmission of LMP1 and associated proteins (2, 3).

Recently, LMP1 has been found to regulate PD-L1 expression at the cellular level through STAT3, AP-1, and NF-κB pathways (34). Cytokines such as tumor necrosis factor-α (TNF-α), interferon-γ (IFN-γ), and interleukin 4 (IL-4) upregulate PD-L1 expression through the STAT and NF-κB pathways (43–45). IFN-γ also has been shown to have a synergetic effect with LMP1 in up-regulating PD-L1 in NPC (34). PD-L1 upregulation has been implicated in numerous roles with regards to immune suppression (28, 43, 44). Constitutive expression of PD-L1 on tumor cells is an innate immunity mechanism activated by several pro-survival pathways: MAPK (45), ATM/ATR/Chk1 (46), JAK/STAT (43), and PI3/AKT(44); or by genetic aberrations such as CDK5 disruption (47), loss of PTEN (48), or truncation of the 3’–UTR that stabilizes an increase in PD-L1 transcripts (49). However, constitutive expression of PD-L1 is far less common than IFN induced PD-L1 expression. PD-L1 is expressed in various cancers: gastric cancer (50), melanoma (51), nasopharyngeal carcinoma (52), post-transplant lymphoproliferative disorder (53), non-small cell lung cancer (54), pancreatic cancer (55), lymphomas (56), bladder cancer (57), kidney cancer (58), breast cancer (59), and Hodgkin lymphoma (60). Other than EBV, multiple human herpes virus infections have also been found to result in increased PD-L1 expression such as herpes simplex virus (61), Varicella zoster virus (62), Cytomegalovirus (63), Human herpesvirus 6 (64), and Kaposi’s sarcoma-associated herpesvirus (65). Interestingly, in addition to human herpes viruses, infection with human immunodeficiency virus (HIV), influenza virus, Ebola virus, and adenovirus also leads to elevated PD-L1 expression (66), (67). Altogether, the wide array of diseases viral and non-viral, that are associated with increased PD-L1 expression emphasizes the need for elucidating the mechanism of pathway regulation, activation, and localization of PD-L1.

In immune competent hosts, EBV efficiently infects B lymphocytes, this initiates a strong cytotoxic T cell response against lytic and latent antigens, leading to the expansion of EBV-transformed B cells. To establish and maintain latency, EBV has adopted many immune evasion mechanisms. To further understand the function of PD-L1 in EBV related cancers in response to EBV viral oncogene LMP1, we characterized the subpopulations of extracellular vesicles expressing PD-L1 following LMP1 expression. In our experiment we determined PD-L1 enrichment in MVs in response to LMP1 levels in both a high LMP1 expressing cell line construct (HK1 GFP-tagged inducible LMP1) and by a LMP1 expressing cell line (HNE-1 pBabe LMP1) which constitutively expresses lower levels of LMP1, similar to levels found in EBV infected cells. It was observed that LMP1 and PD-L1 accumulated in a subpopulation of EVs called MVs. Additionally, we determined that the upregulation of PD-L1 in the MV subpopulation was in response to EBV LMP1. MVs from NPC cells were confirmed though size, density and the detection of MV markers. Flow cytometry analyses indicated an approximately 10-fold increase in PD-L1 associated MVs from the NPC cell line expressing LMP1 compared to the wild type. LMP1 upregulated the cellular expression of PD-L1 at the transcriptional level; however interestingly, PD-L1 mRNAs were detected in MVs. PD-L1 is post-transcriptionally regulated in an indirect way through TNFα-activated NF-κB (68). LMP1 is a CD40 receptor mimic that constitutively activates NF-κB (69, 70). This depicts a mechanism in which PD-L1 is possibly being upregulated at the transcriptional level in MVs through the activation of NF-κB by LMP1.

Moreover, we demonstrate that LMP1 expression in NPC cells results in total elevated levels of EV secretion and enhanced packaging of proteins within the MV. One possible explanation for the increase in protein per MV may be that EVs released from virally infected cells can package more proteins involved in suppressing the immune response toward the virus and consequently enhancing viral spread. LMP1 also appears to play a role in enhancing PD-L1 surface exposure in MVs. Enhanced PD-L1 surface levels could depict the mechanism in which LMP1 contributes to immune suppression by enhancing PD-L1-PD-1 interaction and T cell inactivation. LMP1 and PD-L1 also co-migrate within the same density gradient, and immuno-isolation of this MV subpopulation confirmed the presence of LMP1 in PD-L1 containing vesicles from an LMP1 expressing NPC cell line. EGFR was also immuno-isolated within the same MV with LMP1 and PD-L1. Recently, EGFR has been found to play a vital role in upregulating PD-L1 once activated, mediating immune escape in non-small cell lung cancer (71). LMP1 signaling occurs through localization in lipid rafts and by recruiting key signaling components to internal lipid raft containing membranes (35, 72). Our observations indicate that both LMP1 and PD-L1 colocalize perinuclearly within the cell and are enriched in lipid rafts which suggests a similar signaling platform between these two proteins, bringing together numerous signaling components that drive immune evasion mechanisms, and facilitates their interaction. These findings lead us to conclude that LMP1 might have different roles in different subpopulations of EVs, and that LMP1 modified MVs may act as an instrumental component contributing to immune suppression and evasion of EBV viral infected cells, by the enhanced packaging of proteins into these MVs, with one of them being PD-L1.

Altogether, the above data suggests that EBV+ related diseases are largely dependent on PD-L1 for disease progression and possibly immune protection, and identifies a more specific target for treating these diseases. Therefore, any immunotherapy treatments targeted at disrupting this pathway could lead to better outcomes for patients suffering from EBV related diseases. One form of treatment for cancers that induce PD-L1 expression is immunotherapy which serves to boost the immune antitumor response and has fewer side effects than alternative cancer therapies, such as chemotherapy, surgery, and radiation. FDA approved antibodies pembrolizumab (73) and nivolumab (74) that target PD-1 and atezolizumab (75) that targets PD-L1, improve survival against Hodgkin’s lymphoma (73, 75). Anti-PD-1 antibody is effective in several other cancers, such as glioblastoma (76), and metastatic melanoma (28). In addition to cancer, anti-PD-1/PD-L1 treatment has also been considered as a possible treatment for patients with dementia and Alzheimer’s (77). Moreover, monoclonal antibodies targeting PD-1 or PD-L1 are now in phase I-II trials as a therapeutic treatment for patients with metastatic/recurrent NPC (78). Using anti-PD-1 or anti-PD-L1 ligands alone is usually insufficient. Unfortunately, using several immune blockade therapies increases the patient’s likelihood of experiencing unwanted side effects (79).

Now that EBV’s role in the regulation of PD-L1 has been established, exploring other viral functions that might be inducing MV associated PD-L1, such as EBV microRNAs, EBNA-1/2, and LMP1/2A should be investigated. Targeting both LMP1 and PD-L1 expressing MVs and small EVs in EBV+ tumors might lead to a synergistic inhibitory effect on tumor growth and metastasis. Identifying post-treatment levels of MV levels of PD-L1 with anti-PD-L1 blockade therapy could help distinguish patients who respond to this specific type of therapy early on. Tumor genomic profiling is therefore necessary in order to prescribe the most effective treatment that will confer the best results with the least amount of toxicity and side effects to the patient. Additionally, determining the best way to overcome PD-L1 blockade resistance and detect/target MV associated PD-L1 are the necessary next steps to take in the PD-L1 therapeutic research domain. These results highlight the crucial role that MV associated PD-L1 might be playing in creating an immunosuppressive environment that cancers and possibly other diseases exploit. As more research on MV associated PD-L1 in EBV related cancers is conducted, its vital role in facilitating tumor progression can be elucidated.

The results from this research contribute to the EV field and PD-L1 immune-targeted therapies in several ways. Determining that MVs derived from EBV+ cell lines express PD-L1, identifies a new therapeutic target in immune blockade therapy. Consequently, determining whether PD-L1 localization is based on cell line, or on levels of LMP1 will help clarify the mechanisms that promote immune suppression since PD-L1 surface exposure is necessary for T cell inactivation. The next steps include determining the immunosuppressive functions of MV surface expressed PD-L1 in EBV related cancers and determining if they differ from PD-L1 function within small EVs. This immunosuppressive contribution could equate to exosomal levels of PD-L1 or may play a more vital role in immune suppression. In conclusion, we expect that our findings will begin to clarify the mechanism of LMP1-mediated enhanced packaging of PD-L1 into MVs and the functions of LMP1-modified MVs. In terms of clinical applications, it could also serve as a potential therapeutic target or diagnostic biomarker for identifying whether or not the patient will respond well to immunotherapy treatment and predict tumor responses.

## Materials and Methods

### Retrovirus production

Retrovirus particles for transduction and stable cell generation were produced in HNE-1 cells following JetPrime transfection of expression plasmids (pBabe neo, pBabe-HA-LMP1 neo) and packaging plasmids pMD2.G (Addgene; number 12259; a gift from Didier Trono) and PSPAX2 (Addgene; number 12260; a gift from Didier Trono) according to the manufacturer’s instructions (Polyplus). Medium was collected at 48, 72, and 96 h post-transfection, centrifuged for 10 min at 1,000 × *g*, filtered through a 0.45-μm filter, and frozen at −80°C until use. Stable HNE1 cell lines expressing pBabe-HA-LMP1 were created by retrovirus transduction as described above and were selected and maintained in 1 mg/ml of G418 sulfate.

### Generation of GFP-LMP1-inducible cells

HK1 cells that stably express GFP-LMP1 under control of a tetracycline-inducible promoter were created by first being transduced with lentivirus particles containing pLenti CMV TetR BLAST (Addgene; 17492) as previously described in (18). Stable cells were selected with medium containing 10 μg/ml of blasticidin (InvivoGen; ant-bl-1) and then transduced with retrovirus particles containing pQCXP GFP-LMP1. Doubly stable cells were selected with medium supplemented with blasticidin (10 μg/ml) and puromycin (2 μg/ml) for 2 weeks. LMP1 expression was induced for 24 h with the addition of doxycycline to a final concentration of 1 μg/ml or 0.5 μg/ml.

### Cell culture

HNE1 cells were cultured in 1:1 mixture of Dulbecco modified Eagle medium (DMEM; Lonza; 12-604Q) and Ham’s F-12. HK1 (a gift from George Tsao, Hong Kong University). All cell culture Medium was supplemented with 10% fetal bovine serum (FBS; Seradigm; 1400-500), 2 mM l-glutamine (Corning; 25-005-CI), 100 IU of penicillin-streptomycin (Corning; 30-002-CI), and 100 μg/ml:0.25 μg/ml antibiotic/antimycotic (Corning; 30-002-CI). The cells were maintained at 37°C with 5% CO_2_.

### Transient Transfection

HK1 grown in 100 mm plates were transfected with 5 μg of RFP-LMP1 or GFP-PD-L1 plasmid using Lipofectamine 3000 transfection kit (Invitrogen, L3000015). Lipofectamine reagents were diluted in Opti-MEM medium (Gibco, 31,985–070) and added according to manufacturer’s instructions. Twenty-four hours after transfection, cells were harvested and lysed in RIPA buffer, as previously described (25, 47). Cell-conditioned medium was harvested for extracellular vesicle enrichment.

### Immunofluorescence assay

Cells were stained as previously described (47). The coverslips were coated with poly-L-lysine (Sigma; P1955) as according to manufacturer’s instructions, then cells were seeded. The next day, cells had GFP-LMP1 induced using 0.5 μg/ml of doxycycline. The next day, cells were fixed in 100% ice-cold methanol or 4% paraformaldehyde for 10 minutes. Cells were then washed again with PBS 3 times before being permeabilized with 0.2% Triton-X in PBS. Cells were then placed in a blocking buffer (5% goat serum in PBS with 0.2% Tween [PBST]) for 30 minutes. Next, cells were stained with primary antibodies in blocking buffer PD-L1 D8T4X (1:200) for 2 hours. Cells were washed 3 times in PBST then placed in secondary antibody rabbit IgG (A48254; Thermo Fisher) diluted in the blocking buffer for 1 hour. Cells were washed 3 times with PBST before being incubated with Red DNA stain (52406; ENZO) diluted in PBS for 10 minutes. Cells were washed 2 times with PBS, and once with water before being mounted on a glass slide with mounting medium (4% propyl gallate, 90% glycerol in PBS) for confocal microscopy imaging. Confocal images were taken using a Zeiss LSM 880 microscope and processed using Zen 2.1 Black software.

### Extracellular-vesicle enrichment

Extracellular vesicles were harvested from conditioned cell culture media by differential centrifugation and polyethylene glycol (PEG) precipitation as previously described (80),(81). Briefly, apoptotic bodies/cell debris (2K), microvesicles (10K), and small EVs (100K) were harvested from cell-conditioned media by ultra-centrifugation at 2,000 RPM for 10 min, 10,000 RPM for 30 min, and by the ExtraPEG method respectively.

### Protease Protection Experiment

After the washing step the MV pellet is resuspended in a 1:1 ratio of 0.25% trypsin and particle free PBS and left at 37°C for 30 minutes. This cleaves the surface membrane of the EV sample. Antibodies: (PD-L1 E1J2J; Cell Signaling) for extracellular domain of PD-L1, and (PD-L1 E1L3N; Cell Signaling) was used for endogenous levels of PD-L1. Sample is then resuspended in lysis buffer for immunoblot analysis.

### Purification of membrane microdomains

Lipid raft (LR) microdomains were harvested as previously described (82).

### Iodixanol density gradient

EVs from HK1 GFP-LMP1-inducible cells and HNE1-pBabe-HA-LMP1 cells harvested by centrifugation for 10,000 x g for 30 min were further purified on a density gradient, according to the method of Kowal et al. (24). Briefly, EV pellets following the centrifugation step were re-suspended in 1.5 ml of 0.25 M sucrose buffer (10 mM Tris [pH 7.4]). A 60% stock (wt/vol) Optiprep (Sigma, D1556) solution was added 1:1 to EV suspensions and transferred to an MLS-50 rotor tube (Beckman; 344057). Iodixanol stock was then diluted in 0.25 M sucrose-Tris buffer to make 10% and 20% iodixanol solutions, and then 1.3 ml of 20% iodixanol and 1.2 ml of 10% iodixanol solutions were carefully layered on top of EV suspensions. Gradients were centrifuged for 90 min at maximum MLS-50 rotor speed (268,000 × *g*) and separated by collection of 490 μl fractions from the top of the gradient. Densities were measured with a refractometer (Refracto 30PX). Individual fractions were washed with PBS and re-pelleted by ultracentrifugation in an SW41 Ti rotor at 100,000 × *g* for 2 h. Pellets were resuspended in particle-free PBS for NTA or strong urea-containing lysis buffer for immunoblot analysis.

### Immunoblot analysis

To prepare cell lysates, cells were washed and then scraped into cold PBS and collected by centrifugation at 1,000 × *g* for lysis in radio immune precipitation assay (RIPA) buffer (20 mM Tris-HCl, 50 mM NaCl, 1% NP-40, 0.1% SDS, 0.5% deoxycholate) with 1X proteinase inhibitor. EV samples were harvested as described above. To prepare all cell lysates, and EV lysates run under reducing conditions for SDS-PAGE, additional sample buffer (5×) also containing 0.2 M dithiothreitol (DTT) was added to samples. Lysates were boiled for 5 min. An equal protein concentration or equal volume of cell and EV lysate was run in an SDS 10% polyacrylamide gel and subsequently transferred onto a nitrocellulose membrane (GE Healthcare). Total protein was measured by Ponceau S stain. Blots were blocked with 5% (wt./vol.) nonfat dry milk powder in Tris-buffered saline with Tween 20 (TBS-T). Blots were probed with primary antibodies against the following: PD-L1 (E1L3N; Cell Signaling), Syntenin-1 (S-31; Santa Cruz), HSC70 (B-6; Santa Cruz), Caveolin-1 (D46G3; Cell Signaling), CD63 (TS63; Abcam), LMP1 (CS1-4; Dako), CD81 (B399; GeneTex), CD9 (CBL162; Millipore), CD71 (D7G9X; Cell Signaling), EGFR (SC-03; Santa Cruz). Blots were subsequently probed with the following horseradish peroxidase (HRP)-conjugated secondary antibodies: rabbit anti-mouse IgG (Genetex; 26728), or anti-mouse kappa light chain (H139-52.1; Abcam). Blots were imaged using an ImageQuant LAS4000 (General Electric) and processed with ImageQuant TL v8.1.0.0 software, Adobe Photoshop CS6, and Adobe illustrator.

### Nanoparticle tracking

Nanoparticle tracking analysis (NTA), a technology used to quantify nanoparticle concentrations and sizes consistent with EV populations, was used to determine increases in vesicle secretion and confirm size range of EV pellets following PD-L1 expression in cells. Following EV enrichment, vesicles were resuspended in particle-free PBS for quantitation using a Malvern NanoSight LM10 instrument as previously described in detail (83)(81). The camera level was set to 13, and the threshold was maintained at 3 for all samples. The quantity of particles measured by NTA was normalized to the number of live cells counted at the time of harvest to generate a measure of the number of EVs secreted per cell. For nanoparticle tracking experiments, live cells were counted with an automated cell counter (Cellometer Vision, software version 2.1.4.2; Nexcelom Biosciences) at the time of harvest by staining with a 0.2% final concentration of trypan blue (Sigma; T8154) in phosphate-buffered saline (PBS) in a 1:1 ratio. Relative vesicle secretion was obtained by normalization of EV levels to control cells in each experiment.

### Transmission electron microscopy

Following enrichment of EVs by the ExtraPEG method, pellets were resuspended in 100 μl of particle-free PBS for electron microscopy imaging. Samples were prepared as described by Lässer et al. (84) and visualized on an FEI CM120 transmission electron microscope. For immunogold labelling, EV pellet was resuspended in anti-PD-L1 antibody (D8T4X; Cell Signaling) (1:60) in blocking buffer (1% BSA in PBS) for 2 hours. The EV suspensions were applied to formvar/carbon-coated nickel grids 200 mesh for 3 min. Then grids were washed with 5 separate drops (50 μl, 10 min per drop) of PBS with 0.1% BSA. Transferred a drop of the secondary antibody conjugated to 10 nm gold particle (Goat F(ab’)2 Anti-Rabbit IgG H&L (10nm Gold; ab39601) (1:100 in PBS with 0.1%BSA) to grids for 1 hour. Repeat washing steps. Next wash with 2 separate drops of fresh MilliQ. Finally, negative stain with uranyl acetate for 1 min.

### Live cell imaging

HNE-1 cells were seeded into 35-mm glass well plates (Greiner Bio-One; number 627860), co-transfected 24 h later with 1 μg of GFP tagged PD-L1 and RFP tagged LMP1, and imaged 24 h post transfection using a Zeiss LSM 880 microscope with a live-cell imaging chamber. Hoechst 33342 nuclear staining (5 μg/ml; Thermo Scientific; number 62249) was added 15 min prior to imaging. Confocal images were taken and processed using Zen 2.1 Black software.

### Immunoprecipitation

The MV sample obtained after centrifugation was resuspended in 1%BSA in PBS. 100 μl of magnetic G beads were washed 3 times with (0.1%BSA in PBS). Rabbit (DA1E) mAb IgG XP® Isotype Control or PD-L1 (Extracellular Domain Specific) (D8T4X) antibodies were diluted in PBS with 0.1% BSA and then mixed with 1.0 μg magnetic beads. The antibodies and magnetic beads were incubated with rotation for 20 min at room temperature. This was followed by three washes with 0.1% BSA in PBS. The Microvesicle sample was then added to the beads and left in the cold room for 3 hours with rotation. This was followed by another three washes with 0.1% BSA in PBS. Resuspended in 2x reducing buffer and then heat for 7 min at 90 °C before running on gel for immunoblot analysis.

### Flow cytometry

Microvesicles from HK1 or HK1-LMP1 cell were first blocked with flow buffer (1% BSA in PBS) stained with either isotype control Ig or anti-PDL1 at 2g/mL for one hour at 4 °C. After wash with flow buffer, samples were stained with PE conjugated anti-rabbit antibody at 4 °C for 30 mins. Wash to remove unbound antibodies, samples were run on FACSC onto flow cytometer (BD) in Translational lab College of Medicine FSU. The data were analyzed by FACSDiva software (BD).

### RNA Isolation and Reverse Transcription

Total RNA of cell or EV samples were isolated by Trizol reagent (ThermoFisher, 15596018) and quantified by nanodrop. Less than 1 μg of total RNA was used for reverse transcription by qScript cDNA SuperMix (Quantabio, 95048). CDNAs were store in −20°C until further use.

### Quantitative Real-Time PCR and Data analysis

Standard 3-step cycles protocol (40 cycles of 95°C for 5 s, 60°C for 10 s, 72°C for 20 s) was used in all qPCR reactions. PerfeCTa SYBR Green FastMix (Quantabio, 95072-012), assay primers (250 nM) and cDNA (1.0 μL) of cell or EV were prepared in 20 μL reaction and run on CFX96 qPCR machine (Bio-Rad). Gene expression level were first normalized to housekeeping gene GAPDH and then calculated with ΔΔCt method. qPCR primer sequence:

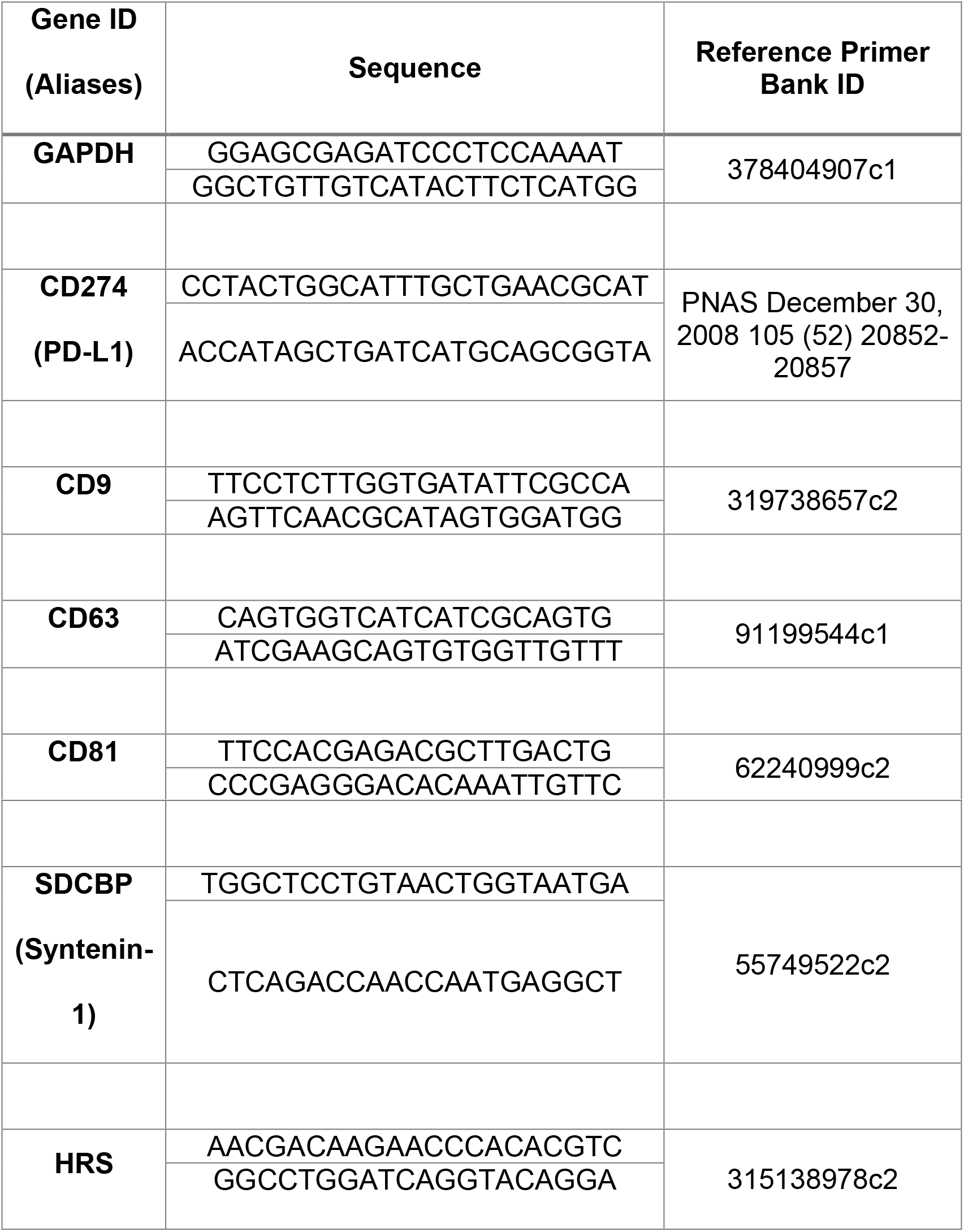

### Data and statistics analysis

Statistical analysis was performed using the GraphPad Prism 8 (GraphPad Software, San Diego, CA) or Microsoft Excel with a significant threshold of p≤0.05. Pearson correlation coefficient (PCC) was determined using an ImageJ colocalization plugin, with a minimum of 10 cells analyzed. The ImageJ software (NIH) was used to calculate the CTCF by applying this formula: CTCF = integrated density (indicated by the software)-(Area of selected cell x Mean fluorescence of background readings), as previously described (85). Statistical significance of results was evaluated by unpaired t test with Welch’s correction. Figures were constructed using Adobe illustrator.

## Acknowledgments

We would like to thank Dingani Nkosi and Sara York with their help on experiments and useful discussions. Special thanks to Yah Xin at the National High Magnetic Field Laboratory for help obtaining electron microscopy images and the FSU College of Medicine Confocal Microscopy Laboratory. This research was supported by a grant from the National Cancer Institute of the National Institutes of Health (RO1CA204621) awarded to D.G.M.

## Author Contributions

Conceptualization, M.A. and D.G.M.; data curation, M.A. and L.S.; funding acquisition, D.G.M.; investigation, M.A. and L.S. methodology, M.A., L.S. and D.G.M; project administration, D.G.M.; software, M.A. and L.S..; supervision, D.G.M.; writing–original draft, M.A.; writing—review and editing, D.G.M. and L.S. All authors have read and agreed to the published version of the manuscript.

## Conflict of Interest

The authors report no conflicts of interest. The funders had no role in the design of the study; in the collection, analyses, or interpretation of data; in the writing of the manuscript, or in the decision to publish the results.

